# Mechanism of HIV-1 Capsid Rupture and Uncoating by Reverse Transcription

**DOI:** 10.64898/2026.04.17.719300

**Authors:** Kuntal Ghosh, Manish Gupta, Gregory A. Voth

## Abstract

One of the key events in the HIV-1 life cycle is reverse transcription, during which single-stranded viral RNA (ssRNA) is converted into double-stranded DNA (dsDNA). This process occurs inside the mature virus capsid and, once it reaches a critical threshold, drives capsid rupture. This uncoating is essential for infection because it releases viral genetic material into the host cell nucleus. Despite its importance, many mechanistic details of this process remain to be fully understood. To address this gap, we develop a multiscale computational method for simulating reverse transcription inside the capsid, termed Coarse-Grained Kinetic Monte Carlo (CG-KMC). CG-KMC stochastically adds deoxynucleotide triphosphates (dNTPs) to the coarse-grained RNA model, enabling stepwise growth of DNA inside the HIV-1 capsid. We implement this method within an integrative coarse-grained framework that combines a “bottom-up” capsid model with a “top-down” representation of the viral RNA/DNA genome. Our simulations phenomenologically capture and predict diverse capsid rupture pathways during reverse transcription. The resulting ruptured structures closely match previously identified cryo-ET images. We further perform an extensive analysis of the rupture process, examining its mechanistic and kinetic aspects as well as the role of capsid–DNA interactions. Our findings illuminate how different capsid–DNA conditions give rise to distinct rupture pathways, which differ from ruptures due to simple outward pressure expansion models from within the capsid.

**Significance Statement:** Reverse transcription (RT) is a critical step in the life cycle of HIV-1, the causative agent of the AIDS pandemic. During RT, single-stranded RNA (ssRNA), encapsulated inside a mature capsid, is converted into double-stranded DNA (dsDNA). The rigidity of dsDNA generates increased internal pressure inside the capsid which, beyond a critical threshold, drives capsid uncoating, releasing the viral genome into the infected host cell. However, key mechanistic and kinetic aspects of this process remain unresolved. Here, using coarse-grained simulations, we elucidate the mechanistic basis of RT-induced capsid uncoating and quantify the role of capsid-genome interactions. Our results demonstrate that tuning these interactions may provide a potential antiviral strategy to inhibit infectivity by targeting RT.

## Introduction

The HIV-1 life cycle comprises a sequence of tightly regulated steps, from particle assembly and protein biosynthesis to the integration of the viral genome into the host DNA for infection (1). Among these, the process of reverse transcription (RT) – the conversion of flexible viral single-stranded RNA (ssRNA) genome into stiff double-stranded DNA (dsDNA) within the mature capsid – is particularly critical yet still not fully understood (2). Experimental studies (3–5) have probed RT; however, a unified consensus on its timing and the sequence of accompanying events has yet to emerge. It is also fairly well established that RT within the viral capsid is intricately linked to the capsid uncoating (6–11), involving its partial or complete disassembly. However, the precise timeline of RT relative to capsid destabilization and nuclear entry remains unresolved. As RT proceeds, the encapsulated ssRNA gradually transforms into dsDNA, exerting considerable but not homogeneous mechanical force on the capsid interior wall that contributes to capsid rupture.

A mature HIV-1 capsid is composed of capsid (CA) protein dimers that self-assemble into a range of morphologies (12, 13). In its canonical mature form, the capsid adopts a closed fullerene-like, cone-shaped shell built from approximately 250 hexamers and precisely 12 pentamers (14, 15). This shell encloses the viral ssRNA genome together with several essential enzymes, including protease, reverse transcriptase, and integrase (16, 17). It was originally thought that RT is initiated and completed in the cytoplasm (2, 4), even before the capsid reaches the nuclear pore complex (NPC), which characterizes the boundary between the nucleus and the cytoplasm. More recent studies, however, suggest that RT proceeds as the capsid is docking at or traversing the NPC, and may even be completed inside the nucleus (6, 18). These observations underscore a critical gap: although the biochemical stages of RT are well characterized, their temporal coordination with capsid structural transitions during and after nuclear entry remains poorly defined. Furthermore, the detailed mechanistic pathways leading to the capsid uncoating process are not well understood. To address these challenges, we developed and deployed in this work large-scale coarse-grained (CG) models of the RT process inside a mature HIV-1 capsid. Coarse-graining (19, 20) involves constructing a simplified representation of a much larger atomistically resolved system, with the goal of retaining its primary physical features and effective interactions at a fraction of the computational cost. All-atom and very large scale molecular dynamics (AA MD) has been used previously (15, 21, 22) to examine the HIV-1 capsid shell. One of these papers (22) presented a more detailed model that revealed that when the CA lattice-bound anion IP6 and a model for the genome package were included in the AA MD simulation model, the capsid shell developed distinct inhomogeneous strain (lattice compression and expansion) patterns. This result was also suggestive of non-uniform capsid rupture behavior as the internal RT process proceeds.

On the other hand, simulating the complete RT process within a capsid with AA MD is computationally intractable. However, in the past few years CG-MD has had considerable success in modeling HIV-1 capsid maturation and host interactions (13, 23–27), as well as simulating the entry of the capsid into the nuclear pore complex (NPC) (28, 29). Building on these developments, in this work, we develop an integrative CG model of a “bottom-up” HIV-1 capsid (derived from the AA interactions) with a top-down genome model (encompassing both RNA and DNA). We develop and implement a stochastic Kinetic Monte Carlo (30, 31) (KMC) method at the CG level, which we denote as Coarse-grained Kinetic Monte Carlo (CG-KMC), to model RT and subsequent capsid rupture and uncoating. We show that our model accurately captures capsid rupture as RT progresses, in striking agreement with previously reported experimental evidence as imaged by cryo-electron tomography (5). We also investigate and characterize the impact of the varying capsid-DNA interactions on the rupture pathways.

## Results

### Capsids stay intact during initial stages of reverse transcription

A condensed “ball” of RNA was embedded inside the cone-shaped HIV-1 capsid (**Figure 1A**). In concordance with experiments (32), capsid uncoating does not begin instantaneously in the presence of dNTPs. Several critical processes involving enzymes such as reverse transcriptase (33–35) (incorporated implicitly) facilitate the growth of RT products from single-stranded copies of RNA. Stages I and II (**Figures 1B-C**; described in the Materials and Methods section) depict the growth of the RNA-DNA hybrid, formed stochastically with the addition of one-site CG dNTPs to the RNA beads, and the subsequent cleavage of the hybrid. The reader is directed to the SI for a discussion of the polypurine tracts (PPT) (36). As is evident in **Figure 1C**, growth of the RNA-DNA hybrid takes place inside the capsid, without leading to distortions in the capsid lattice (37). It is also important to note that both the initial ssRNA and the RNA-DNA hybrid strands are modeled to be sufficiently flexible, thus allowing them to “coil up” as they keep growing as a result of the CG-KMC algorithm. Even with respect to Stage III in our model (**Figure 1D**; described in the Materials and Methods section) which generates dsDNA, there is no rupture at the beginning. This can be corroborated well by the experimental notion that the growth of dsDNA leads to rupture only after a certain time interval (∼ 6-10 hours) (5, 32), and after attaining a certain length of the genome (∼ 3.5 kb or about one-third of the genome size) (32).

**Figure 1:**
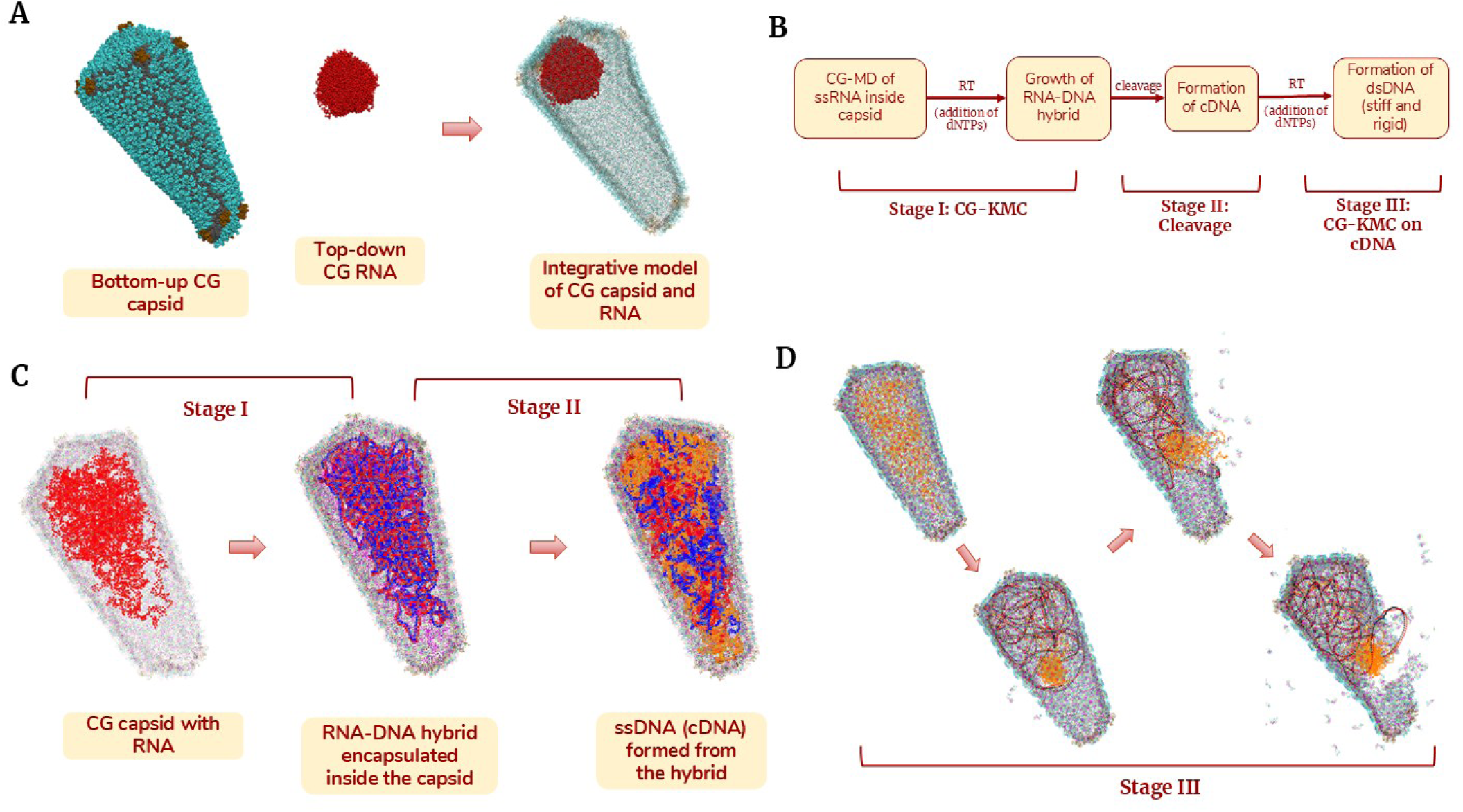
Overview of the integrative CG model and CG-KMC scheme: **(A)** Left: A CG capsid model constructed systematically using a “bottom-up” approach from underlying atomistic trajectories (NTD of hexamers in cyan, NTD of pentamers in ochre, CTD of both in gray); Middle: A highly condensed “ball” of RNA (red), designed in a “top-down” fashion; Right: The RNA is embedded inside the capsid to generate the initial configuration of the capsid-RNA complex, subsequently used for CG-KMC simulation. Pentamers are colored brown in the CG capsid. **(B)** Stages of reverse transcription: CG-MD simulations of dNTP addition to ssRNA (implicitly catalyzed by reverse transcriptase) to form RNA-DNA hybrid. This undergoes cleavage forming cDNA, which again undergoes RT to finally form dsDNA. This entire protocol can be represented by three stages – Stage I (CG-KMC), Stage II (Cleavage) and Stage III (CG-KMC on cDNA). **(C)** Stages I and II of RT: The initial configuration of the capsid with a condensed “ball” of RNA (red) gradually occupies the available volume inside the capsid after a few CG-MD steps. In Stage I, dNTPs stochastically and sequentially bind to the RNA beads using CG-KMC, forming the RNA-DNA hybrid (blue-red complex). Stage II depicts the subsequent degradation of the RNA-DNA hybrid, in a type of “cleavage”, which yields a single-stranded complementary DNA (cDNA) (orange). **(D)** Stage III of RT: Dynamic growth of dsDNA (red-black hybrid) from cDNA leads to eventual capsid rupture. One representative pathway is shown here.

### Capsid starts uncoating as the double-stranded DNA grows

In Stage III of RT (**Figure 1D**), CG-KMC stochastically keeps adding the dNTPs to the single-stranded cDNA, mimicking the growth of dsDNA. As mentioned before and experimentally established, dsDNA is much stiffer (i.e, has a much higher persistence length) than the flexible ssRNA. As dNTPs are sequentially incorporated into the cDNA, a corresponding strand assembles on the template, with sequential bead–bead connections introduced to model the underlying base-pair interactions. To ensure rigidity and stiffness of the DNA, harmonic interactions were used for modeling the base pair interactions. As the dsDNA grows, it progressively occupies the capsid interior, repeatedly impinging on the inner surface of the capsid lattice. This, however, does not lead to rupture in the initial stages. This behavior in the model can be interpreted as the implicit effect of the nucleocapsid (NC) protein, which is experimentally known for condensing the dsDNA (8, 32, 38). However, the amount of NC protein present cannot condense the full length of the dsDNA, at which point the internal mechanical pressure imparted by the dsDNA on the walls of the capsid increases, eventually leading to rupture. At certain genome lengths, the dsDNA appears to exert substantial pressure on the capsid walls, as indicated by a marked reduction in nucleic acid strand mobility. Each CG-MD trajectory samples a distinct conformational state of the cDNA as it undergoes RT, thereby generating heterogeneous patterns of mechanical stress imparted by the dsDNA on the capsid interior, as opposed to a uniform outward pressure. Through this ensemble of trajectories, our model captures the inherently stochastic nature of reverse transcription and capsid uncoating with both efficiency and at least semi-quantitative fidelity.

The pathway depicted in **Figure S1A** shows initial stages of uncoating along the walls of the capsid. Conversely, the pathway depicted in **Figure S1B** reveals stages of uncoating proceeding from the wider and even narrower ends of the capsid. Rupturing at either end of the cone-shaped capsid is typically characterized by detachment of pentamers located in the high-curvature regions. **Figures S2A-B** depict gradual detachment of pentamers from the wider end of the cone-capsid lattice, whereas **Figures S2C-D** depict gradual uncoating near the narrow end, both directly at the tip and through longitudinal propagation of cracks initiated at the midsection of the capsid. In each of these pathways, uncoating is initiated by the formation of small local patches, succeeded by the progression of uncoating along the cracks, leading to much larger patches. Eventually the capsid disintegrates significantly, thus facilitating release of the viral genome. Two representative pathways of capsid rupture, including the dynamic growth of dsDNA from ssRNA, are shown in **Movie S1**.

We also computed the Green-Lagrange strain patterns (39) (**Figure S7)** to highlight the regions of the capsid walls experiencing increased strain, as an indication of the tentative uncoating domains. Here, we show the evolution of strain patterns at CG resolution for an illustrative case. As RT progresses, strain patterns appear in local patches on the capsid walls, encompassing the hexamers. As this local patch expands along the walls, initial stages of uncoating can be seen, with the most strained CG beads getting detached from the lattice. As this progresses, the beads experiencing the most strain get further detached from the walls, eventually leading to partial disassembly. While our group’s previous study (22) examined the strain patterns that could be associated with capsid uncoating, it did not provide a systematic framework for inducing uncoating driven by RT. By contrast, the present work has an explicit dynamic description of RT inside the capsid, and the predicted rupture patterns are in agreement with experimentally determined lattice separation (correlated with the strain) patterns (22).

We also explored capsid rupture patterns as a consequence of expanding an idealized “virtual object” inside the capsid such as a sphere or cone. The reader is directed to the SI for a detailed discussion. This approach does not include an explicit description of the viral genome inside the capsid. **Figure S9** depicts capsid rupture pathways obtained by isotropically expanding a single virtual sphere and two virtual spheres stacked on top of each other within the capsid. As the rupture patterns show, some configurations are not very realistic, and unlike the RT-driven capsid rupture, the virtual object expansion approach appears more deterministic. The growth of the same geometric object leads to identical rupture pathways across multiple replicas. Thus, to obtain a distribution of rupture pathways which CG-KMC naturally produces, one must tune the geometric parameters of each virtual object. This underscores the necessity of using an explicit description of the dsDNA growth inside the capsid, which can effectively drive anisotropic capsid rupture, as done in this work. We emphasize again that the capsid model utilized here is one derived bottom-up from realistic atomic-scale interactions via a rigorous coarse-graining approach based on statistical thermodynamics.

### Impact of capsid-DNA interactions on uncoating pathways

Capsid-DNA interactions must play a critical role in determining the nature of the disassembly pathway. However, not much is known about the exact nature of interactions governing the initial rupturing of the capsid walls due to the growing dsDNA strands. To this end, we systematically carried out an analysis probing into the role of the CA-DNA interactions, and how it governs the nature of capsid rupture. Modified Lennard-Jones (LJ) interactions (Eq 1) and Gaussian interaction potentials (Eq 2) were used to model the CA-DNA interactions. Owing to the nature of that aspect of our model, we explored different attraction parameters in the modified LJ interactions. It must be pointed out that the same set of parameters were used for the CG simulations before the critical genome size is reached, at which point, the attractive potential is switched on (which we call an experimentally motivated “switch”, used to mimic the release of the condensing effect of NC). It is this switch that triggers capsid rupture once the growing dsDNA reaches the experimentally determined critical length—approximately one-third of the full genome (32). We subsequently carried out multiple sets of CG-KMC simulations starting from the initial capsid and RNA model, with three different attractive parameters given by ɛ = 2.0, 3.5 and 5.0 kcal/mol (in Eq 1).

**Figures 2, S3** and **S4** illustrate snapshots of representative uncoating pathways for each of these values, with a visual representation of the growth of dsDNA inside the capsid interior from the cDNA, and subsequent rupture. We also report the fraction of the genome corresponding to a given configuration, denoted as χ, defined as the number of CG beads that have undergone RT divided by the total number of CG beads in the genome (3000 in this case). As RT progresses, dsDNA punctures the wider end of the capsid lattice, prominently looping out (**Figure S3**). Such observations can also be made for ɛ = 3.5 (**Figure 2**) and ɛ = 5.0 (**Figure S4**); however, rupturing is much more pronounced for the latter. As these dsDNA protrusions puncture the capsid walls, strands of cDNA can sometimes be seen leaking out. For a more comprehensive analysis, we look at multiple rupturing pathways as a function of the CA-DNA interactions in **Figures 3, S5** and **S6** (discussed again in a later section on kinetics). In brief, a higher attractive parameter often leads to formation of more local patches and aggressive rupturing, effectively “crumbling” the capsid lattice. The partially disassembled capsid structures, as predicted by our CG models, agree well with tomographic slices of ruptured cores observed 8-10 hours under standard endogenous RT conditions (5).

**Figure 2:**
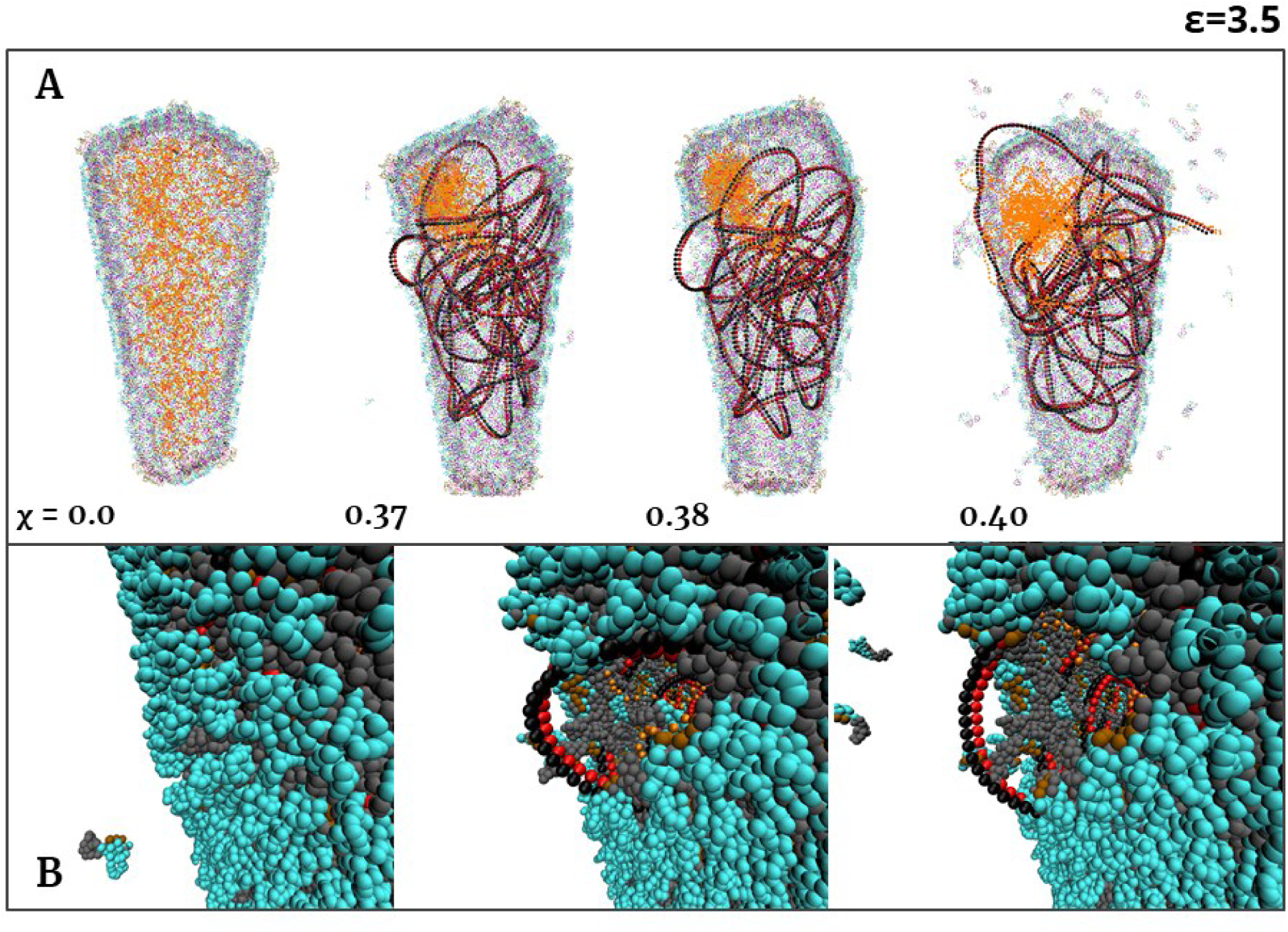
For ɛ = 3.5 kcal/mol, the growth of dsDNA is depicted: **(A)** Snapshots of the capsid enclosing the genome are shown. Initially, the genome is the cDNA (orange). CG-KMC is carried out on the cDNA to form dsDNA (black–red hybrid). As dsDNA grows, rupturing occurs near the midsection of the capsid, and the dsDNA eventually loops out prominently. **(B)** A closer view of the uncoating process. Clear signs of dsDNA protrusion can be seen, accompanied by detachment of capsid hexamers from the lattice. As RT progresses, the capsid walls begin to crumble, releasing loops of dsDNA. Only one representative pathway for ɛ = 3.5 kcal/mol is shown. Here, χ represents the fraction of the total genome size, and the approximate CG timestamp corresponding to a configuration can be given as χ × 3000 × 10^4^CG steps.

**Figure 3:**
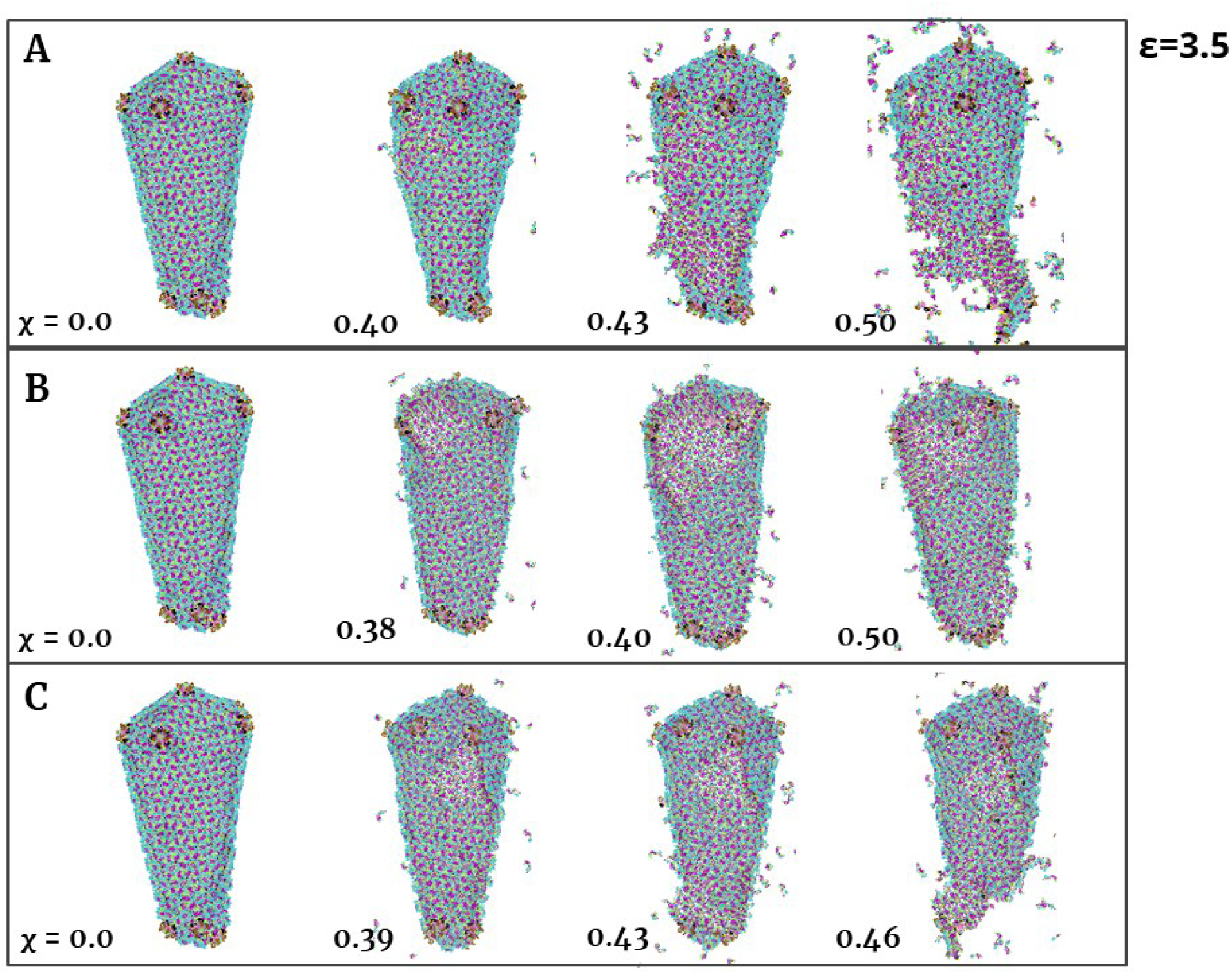
Three representative capsid uncoating pathways for ɛ = 3.5 kcal/mol: **(A)** Cracks first appear in the midsection of the capsid lattice as RT progresses, gradually widening into larger openings that lead to partial disassembly. **(B)** Openings that arise near the wider end of the capsid steadily expand and are accompanied by the detachment of both pentamers and hexamers. **(C)** Cracks originating in the midsection propagate toward the tip of the capsid, ultimately culminating in pronounced disassembly at the tip. Here, χ represents the fraction of the total genome size, and the approximate CG timestamp corresponding to a configuration can be given as χ × 3000 × 10^4^CG steps.

We also present here in **Figure 4** a comparison between representative CG-KMC predicted capsid rupturing patterns with the cryo-ET images of HIV-1 cores during rupture, as reported in Ref. (22). Structures from panels A8, A9, A10 and A12 of Figure 5 and panel 6 of Figure S5 of Ref. (22) are used for comparison. As is evident, CG-KMC predicts cracks near the wider end of the cone-shaped lattice in the regions of higher curvature, which aligns well with the cryo-ET experiments (as in **Figures 4A-B**). More severe rupturing patterns, such as wider holes near the broader end of the cone (**Figure 4C**), or cracks spanning the full length of the capsid (**Figure 4D**) are also predicted.

**Figure 4:**
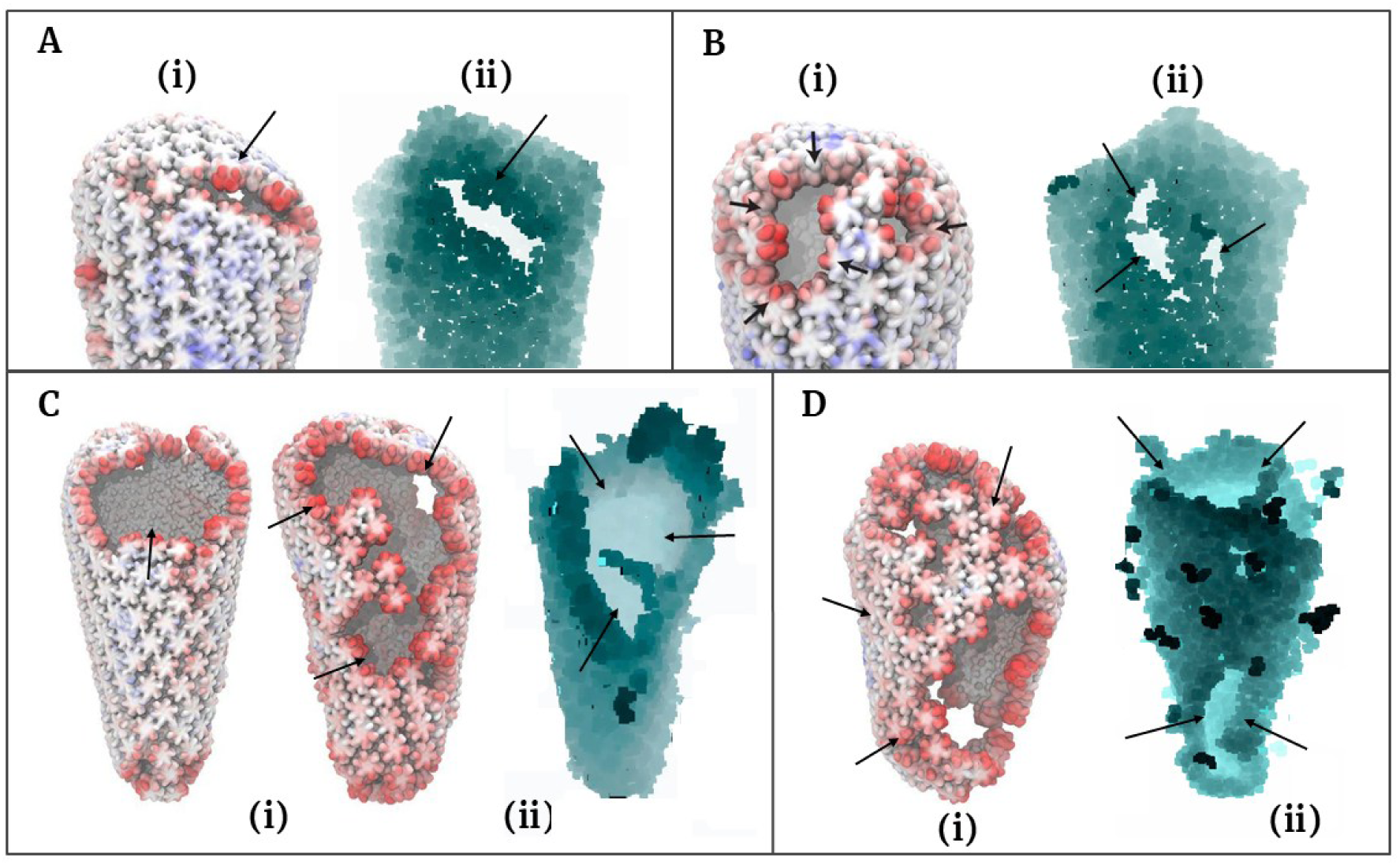
Comparison of CG-KMC predicted capsid rupturing patterns with previously reported cryo-ET images: Subfigures in (i) are cryo-ET images taken from Ref. (22) and those in (ii) show CG-KMC rupturing patterns predicted in this work. Points of rupture are indicated by black arrows. ɛ = 2.0 kcal/mol for A(ii), ɛ = 3.5 kcal/mol for B(ii) and C(ii) and ɛ = 5.0 kcal/mol for D(ii). The groups of dark particles in D(ii) are fragments ruptured from the capsid walls.

**Figure 5:**
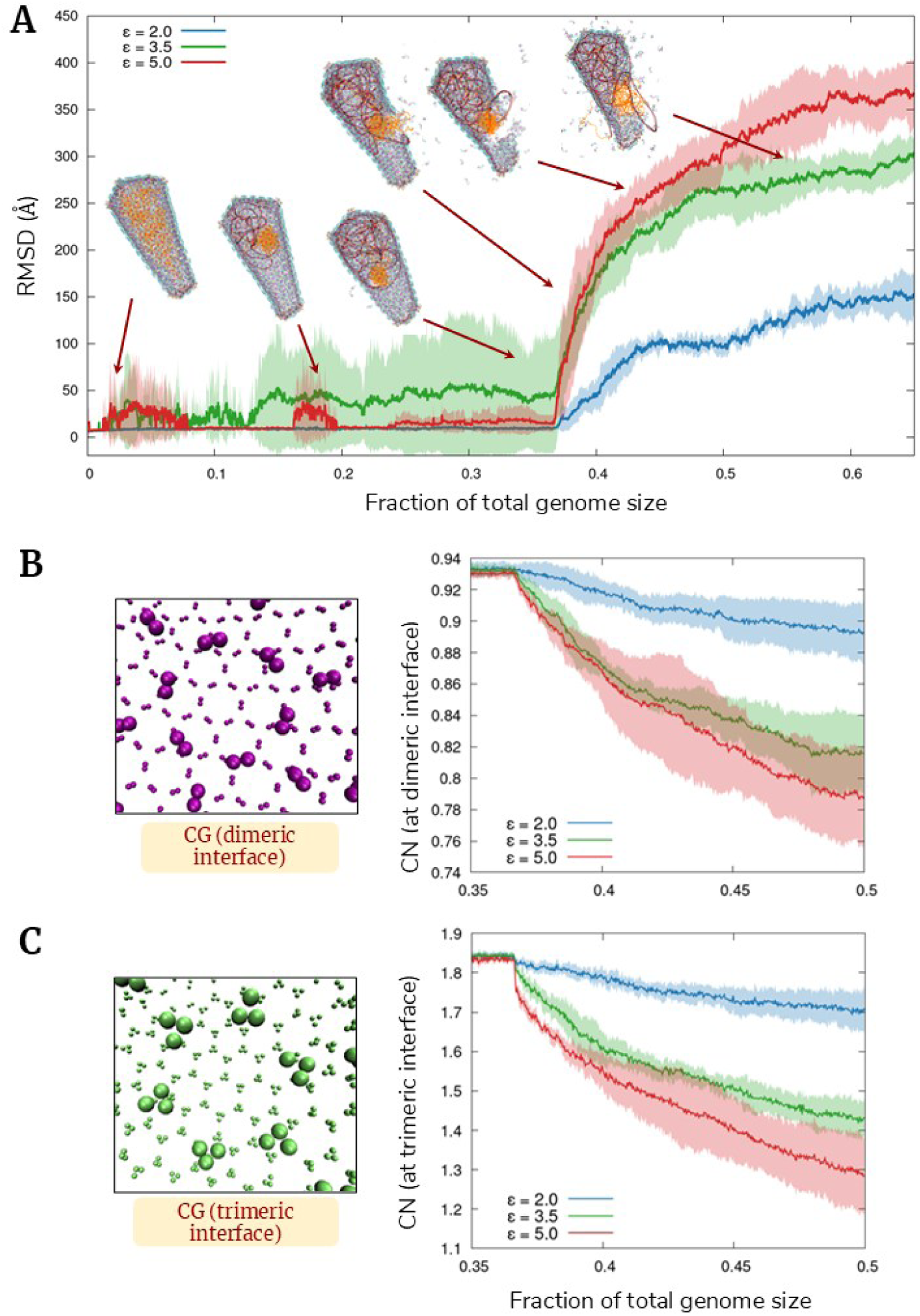
Root-mean-square-deviation (RMSD) and coordination number (CN) analysis: **(A)** RMSD profiles are shown as a function of the fraction of the total genome size (χ) for ɛ = 2.0 (blue), 3.5 (green), and 5.0 (red) kcal/mol. An increase in RMSD indicates a departure from the reference configuration—here, the intact capsid—and thus corresponds to the onset and progression of uncoating. The shaded regions represent the standard deviation of the RMSD values, averaged over three replicas for each ɛ. Regions with elevated RMSD values reflect the inherently stochastic nature of RT. Representative snapshots of capsid rupturing, along with concurrent dsDNA protrusion, are shown at the corresponding χ values at which these events occur. Consistent with experimental observations, the first signs of rupturing appear at roughly one-third of the total genome length. The steepness of the RMSD slope reflects the rate of rupturing progression, which is most pronounced for ɛ = 5.0. As different values of ɛ produce distinct uncoating pathways, only one representative pathway (for ɛ = 3.5) is shown here. **(B)** Left: CG representation of the dimeric interface; Right: evolution of the CN of a CG bead (CN is ∼1 at the dimeric interface). **(C)** Left: CG representation of the trimeric interface; Right: evolution of the CN of a CG bead (CN is ∼2 at the trimeric interface). An increase in the COM–COM distances at these interfaces, or a decrease in the CN, indicates capsid instability—in this case, capsid uncoating. In both cases, as RT progresses, the CN begins to decrease near the critical genome length, with the sharpest decline observed for ε = 5.0 kcal/mol and the most gradual decline for ε = 2.0 kcal/mol.

In addition to a visual analysis of the pathways, we generate root-mean-square-deviation (RMSD) profiles of these pathways as a function of the genome size, with the intact capsid as the reference. From **Figure 5A** it is clearly seen that there is a sharp increase in the RMSD at the critical genome size that corresponds to capsid rupturing. The relative sharpness or “rate” of disassembly progressively increases as the CA-DNA attractive interactions increase. This is consistent with mechanical and biophysical expectations: strengthened CA–DNA interactions increase the effective confinement of the dsDNA, amplifying outward-directed forces on the capsid shell. The resulting rise in internal pressure and elastic strain drives more extensive and pronounced rupture. Notably, the average RMSD for ε = 3.5 kcal/mol is elevated even at early stages, seemingly reflecting the inherently stochastic nature of RT. Before reaching the critical size, the DNA also has greater mobility to explore different regions inside the capsid. At and beyond the critical size, the DNA strands push against the walls much more aggressively, leading to lower strand mobility but increased mechanical pressure. This behavior is also in correspondence with the experimentally observed condensing effect of the NC protein. Thus, despite the absence of explicit NC protein in our CG model, our experiment directed switching scheme implicitly mimics the effect of the NC protein on the dsDNA.

### Kinetic analysis of uncoating events relative to genome size

We also investigated the relative “rate” of rupturing events. While directly correlating uncoating events in real time with CG-MD timesteps is challenging, it is possible to relate them to genome size and changes in CA–DNA interactions. An approximate CG “timestamp” for a given configuration can be estimated as: χ × 3000 × 10^4^ CG steps, where χ is the fraction of the genome size, 3000 is the total number of CG beads in our RNA/DNA model and each KMC move is performed through short CG runs of 10^4^steps (see Materials and Methods). For each of the CA-DNA interactions (ε = 2.0, 3.5 and 5.0 kcal/mol), we qualitatively assess the relative kinetics of uncoating. However, the stochastic nature of the process makes it difficult to generalize precise uncoating timelines. For ε = 2.0 kcal/mol, as mentioned in the previous section, the uncoating is comparatively ‘slower’ than the other cases. Due to the experimentally motivated switch, first stages of rupturing appear at about ∼ one-third the total size of the genome (**Figure 5**). Multiple pathways exhibiting cracks at different regions were predicted (**Figure S5**), including the midsection of the capsid lattice (**Figure S5A**) and near the wider and narrower ends (**Figures S5B-C**).

In all these pathways, cracks propagate slowly and lead to significant rupturing at about half of the genome size (χ = 0.45-0.6), characterized by detachment of pentamers near the wider end and/or ‘opening up’ of the capsid walls at the midsection. It is interesting to note that for certain cases, such as **Figure S5B**, the tip remains intact up to about half of the genome size, and then abruptly uncoats (χ = 0.42-0.53). This can be attributed to the fact that at high curvature regions such as the tip and in local regions of the wide end, the lattice is unable to experience strain, and thus ruptures quickly to release the tension. A similar pattern is observed for the pathway in **Figure 3A** for ε = 3.5 kcal/mol, where the tip ruptures abruptly from a seemingly intact one to a structure with a disassembled lower lattice over a few CG-KMC moves. Even for **Figure 3B**, once a wide patch opens in the lattice, it propagates rapidly toward the wider end of the capsid (χ = 0.38-0.5), further highlighting the role of capsid curvature, as both pentamers and hexamers detach from the lattice. However, the propagation of cracks after the formation of the first patches is the fastest for ε = 5.0 kcal/mol, consistent with our hypothesis (**Figure S6**). For all the pathways analyzed, the initial local patches tend to form near the midsection of the lattice. From there, uncoating proceeds either through further rupture within the midsection itself or through crack propagation toward the tip, ultimately leading to pentamer and hexamer detachment. On an overall note, it can be stated that with an increase in the strength of CA-DNA interactions, the rate of progression of strain and cracks to partial disassembly of the capsid increases drastically after the first signs of uncoating. This is consistent with our RMSD results (**Figure 5A**) and analysis of the interfaces (**Figure 5B-C**).

### Analysis of dimeric and trimeric interfaces during capsid uncoating

While RMSD evolution as a function of the genome size serves as an effective metric for evaluating the overall deviation from the reference structure, a more robust analysis of capsid uncoating is to investigate how the interfaces defining the capsid lattice evolve. A recent paper on the two structural switches in the capsid (40) regulating capsid curvature, host factor binding and capsid entry into the NPC highlighted the structural characteristics of hexameric dimeric and trimeric interfaces influenced by host cofactors such as NUP153. At the CG resolution using a previous mapping scheme (28), the dimeric interface, characterized by V181, W184, M185 and other residues, corresponds to a pair of CG beads. Similarly, the trimeric interface, characterized by I201, L202, L205 and other residues, corresponds to a triplet (**Figures 5B-C**). This corresponds to bead types 37 and 40 for the dimeric and trimeric interfaces respectively in our CG model. Thus, the central geometric ‘cavity’ in the dimeric/trimeric interfaces corresponds to the distance between a pair of CG beads or the area of a triangle formed by a triplet of CG beads. One can monitor the evolution of these interfaces more simply by computing the coordination number (CN) of a representative CG bead at each interface. Thus, for a stable capsid, the CN of a bead at the dimeric and trimeric interfaces is ∼1 and ∼2 respectively, using a 20 A cutoff. As uncoating destabilizes the capsid, it is expected to drastically reduce the CN at each interface, depending on the degree of rupturing. **Figures 5B-C** clearly demonstrates a reduction in the CN at each interface as RT or uncoating progresses. In agreement with the RMSD analysis, increasing the attractive parameters of the CA-DNA interactions leads to a greater degree of rupturing, as the CN demonstrates a much sharper dip at both interfaces due to uncoating. Structurally, this corresponds to a deformation of the hexameric CA-CA units that constitute the capsid lattice. This destabilizes the dimeric and trimeric interfaces between hexamers, progressively compromising lattice integrity and driving partial capsid disassembly.

## Discussion and Concluding Remarks

Investigating the mechanistic and temporal aspects of capsid uncoating driven by RT is highly non-trivial, both experimentally and computationally. From a computational perspective, the primary challenge lies in scale: a mature, solvated HIV-1 capsid undergoing RT would contain > 100 million atoms, making all-atom MD simulations prohibitively expensive, especially given the required long timescales. Experimentally, the intrinsic flexibility and structural heterogeneity of the viral RNA introduce additional complications. Its ability to adopt multiple conformational states makes high-resolution structure determination by x-ray crystallography or cryo-EM difficult (41), thereby limiting the construction of systematic “bottom-up” CG RNA models. Taken together, these challenges necessitate large-scale integrative CG simulations, as implemented in this work, which combine statistical mechanical frameworks with experimentally motivated modeling.

To the best of our knowledge, this work presents the first computational framework that explicitly captures HIV-1 capsid uncoating as a dynamic consequence of RT. By developing and implementing CG-KMC, we successfully simulated the kinetic growth of stiff dsDNA from flexible ssRNA inside the capsid, and, by adding an experimentally motivated switch, used to mimic the release of the condensing effect of NC, we simulated capsid uncoating. Configurations of capsid uncoating predicted by our model are in good agreement with cryo-ET images, thereby validating at least in part the integrative computational approach. It should be noted that our CA–RNA and CA–DNA interactions, as well as the experimentally motivated switch, are necessarily “top-down” or phenomenological due to the lack of high-quality genome structures. In contrast, the capsid itself is modeled bottom-up using a rigorous statistical mechanical framework, which supports our overall approach which is primarily focused on the mechanism of RT-initiated capsid rupture.

We also systematically analyzed capsid rupturing, particularly as a function of different CA-DNA interactions. These CA-DNA interactions might be probed through mutational analysis similar to Ref. (42). Our model predicted distinct predominant pathways corresponding to each interaction—stronger interactions led to more pronounced rupturing. It is important to note that uncoating is a stochastic process, making it remarkable that our model can predict diverse uncoating patterns, ranging from local patches at the capsid midsection to substantial disassembly at the narrow end. Our model effectively captures how dsDNA gradually grows, occupies most of the capsid interior, and, upon reaching a critical length, initiates uncoating, progressively leading to severe disassembly.

We also analyzed the relative timescale of rupture events with respect to genome size. As CG simulations eliminate fast degrees of freedom, CG timesteps cannot be directly equated to real timescales. Thus, although our CG-KMC algorithm integrates Monte Carlo steps with MD to enable sequential conversion of a single-stranded helix to double-stranded DNA, it does not incorporate experimentally determined rate constants, such as dNTP addition to RNA, which are not very well characterized yet.

It is essential to note that uncoating is a complex phenomenon involving the disruption of CA-CA interactions across both hexamers and pentamers. These CA-CA interactions are primarily hydrophobic (15, 43, 44), particularly those at the CTD trimer interface, which play a critical role in capsid assembly and stability. A mature capsid also contains hydrophobic pockets with distinct functions, such as the FG-binding pocket between two monomers (described later). Electrostatic and solvation effects may further influence uncoating (21). Although solvent-free, our CG model incorporates the hydrophobic and electrostatic physics of a mature capsid through a bottom-up approach, effectively capturing uncoating events despite the inherent difficulty of modeling hydrophobic effects accurately by CG models (45).

Another important consideration is the presence of cofactors that facilitate capsid activity, such as nuclear entry and uncoating. One such factor, the IP6 anion, promotes capsid assembly and stability (13, 46, 47). As discussed in the Materials and Methods section, while IP6 is not explicitly included in the CG-MD simulations, it is implicitly present because the bottom-up capsid model was constructed from all-atom trajectories that included a solvated capsid patch and IP6. Moreover, the capsid contains several hydrophobic binding sites, such as the FG-binding pocket between two CA monomers (48). Upon nuclear entry, these FG-binding sites interact with phenylalanine-glycine-rich (FG-rich) domains of intrinsically disordered nucleoporins such NUP98 and NUP153 scattered across the NPC. While our previous study on capsid entry through the NPC included some of these domains (28), such components have not yet been incorporated into the RT model presented here. Other relevant cofactors include CPSF6 and Cyclophilin A (48–50), both of which play important roles in nuclear entry, though their effects on RT remain to be studied. We also aim to investigate the effects of the capsid inhibitor Lenacapavir (LEN) (29, 51–53) on RT driven capsid uncoating. Our overall aim is to construct a model capable of simulating reverse transcription concurrently with nuclear entry through the NPC in the presence of these cofactors. We will also incorporate the effect of explicit reverse transcriptase and capsid binding inhibitors (such as NVP and PF74) into our CG model in the future (32).

In conclusion, this work represents a new advance in modeling reverse transcription–driven capsid uncoating and contributes more broadly to our understanding of the virus capsid properties as well as the HIV-1 life cycle in general.

## Materials and Methods

### Integrative CG capsid and RNA modeling

A bottom-up CG capsid model, used for investigating capsid docking into the nuclear pore (28), was used in this work. Details of the all-atom MD simulations and the subsequent construction of the CG capsid model can be found in the SI of Ref. (28). In brief, a “bottom-up” CG capsid model was constructed from underlying all-atom simulations of three CA hexamers complexed with IP6 and FG peptides. FG peptides play a critical role in regulating CA-NPC interactions, and IP6 is well known for regulating capsid stability (22, 46) and influencing capsid uncoating (5). Thus, IP6 has been implicitly incorporated into our CG model. We also developed and implemented a more phenomenological (or “top-down”) model of the RNA and the CA-RNA/CA-DNA interactions. Two copies of CG single-stranded RNA (ssRNA) are enclosed within the capsid, each minimally representing a ∼ 9 kb genome with 3000 beads. A modified Lennard-Jones interaction with a soft core was used for the interactions between the capsid and DNA:

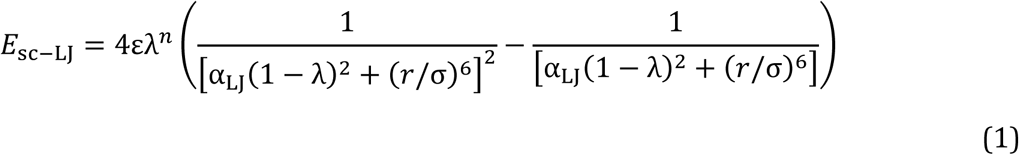

Modeling the CA-RNA and CA-DNA interactions is important as they facilitate capsid uncoating. The CA-DNA interactions were modeled using E_sc-LJ_, and it was the corresponding ε – representing the well depth or strength of attraction – that was varied to investigate how uncoating patterns depend on the CA-DNA interactions. Before a critical threshold, Eq (1) used ε = 0.5 kcal/mol, σ = 12.5 A and λ=0.6. After an experimentally directed “switch” is activated (described later), σ = 7.0 A and λ =0 .6, while ε can be 2.0 or 3.5 or 5.0 kcal/mol (as analyzed extensively in the Results section). In Eq (1) *n*=2 and α_LJ_ = 0.5 were used throughout. dNTPs or deoxynucleotide triphosphates were represented by a one-site CG model. E_sc-LJ_ was also used for modeling CA-dNTP interactions.

Modified Gaussian potentials were used for modeling the electrostatic interactions between the two charged C-terminal CG beads of the capsid with the RNA polymer:

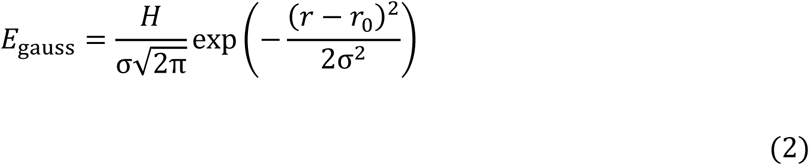

(2)

Where, H = - 4.5 kcal/mol, *r*_0_ = 12.5 A , σ = 1.2 A for the CA-RNA interactions, and H = - 2 kcal/mol for the CA-dNTP interactions.

To prevent CG beads from overlapping with each other, soft repulsive cosine interactions were used:

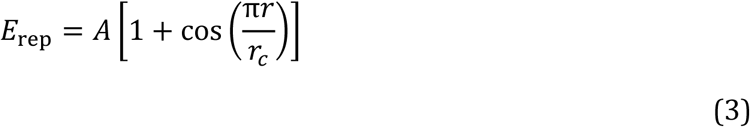

Where, A = 10 kcal/mol for RNA-DNA and 20 kcal/mol for DNA-DNA interactions (between CG beads of different strands), and *r*_*c*_ = 15 A . Eq (3) was also used for the CA-dNTP interactions (A = 1 kcal/mol and *r*_*c*_ = 12.5 A ). All other interaction parameters, and the bonding/angular parameters can be found in the LAMMPS input files.

### CG-KMC for modeling stages of reverse transcription

Reverse transcription is a complex process comprising multiple stages, including several strand transfer events (3, 54, 55). It is computationally intractable to atomistically model such an event as it (i) requires high quality structures of RNA (ii) necessitates the use of very efficient methods capable of modeling bond formation/dissociation. Classical MD with enhanced sampling tools, while being computationally efficient, uses fixed bonding topologies, thus making it impossible to capture reactivity. On another note, *ab initio* or even QM/MM (56) simulations are limited to a few hundred picoseconds. While approaches such as multiscale reactive molecular dynamics (MS-RMD) (57), and reactive methods combined with CG (58, 59) offer a promising approach to enhance chemical reactivity in MD, it is very difficult to efficiently parameterize and devise such methods for large-scale proteins and viruses. To this end, we developed an efficient Coarse-Grained Kinetic Monte Carlo (CG-KMC) approach to model the stages of RT (schematically depicted in **Figure 1B**). Additional variants of KMC schemes have been reported in the literature as well (60).

From a chemical biology perspective, following the first strand transfer (FST) DNA is synthesized along an ssRNA strand. This is followed by minus strand synthesis and a primer generation, followed by the degradation of RNA and plus strand synthesis. Following second strand transfer (SST), a second DNA strand is generated, finally yielding dsDNA (5). As the exact events characterizing RT are very complex, we propose a three-stage approach to model RT: (1) Growth of the RNA-DNA hybrid from ssRNA, (2) Dissociation (or “cleavage”) of the RNA-DNA hybrid, (61) (loosely corresponding to RNA degradation) to form a single-stranded complementary (cDNA) strand, and (3) Formation of dsDNA from cDNA.

The simulations start with a condensed “ball” of RNA (two single-stranded copies) embedded inside the capsid. Reverse transcription progresses by the addition of deoxynucleotide triphosphates (dNTPs) to the RNA. Stage I simulates a stepwise, stochastic, sequential addition of dNTPs, denoted by one CG bead, to one copy of the RNA. Thus, each CG-KMC move permits the formation of a single bond between an RNA bead and a dNTP bead, such that a bond can be formed when the RNA bead in concern is within 20 A of a dNTP bead. The reader is directed to the SI for a mathematical formulation of CG-KMC. Subsequently, corresponding bonds and angles are formed between the dNTP beads adding onto the RNA. This gradually leads to the formation of an RNA-DNA hybrid. It is valuable to note that this RNA-DNA hybrid naturally acquires a helical nature, which qualitatively mimics the helicity of the RNA-DNA hybrid. Following this, the RNA half of the DNA gets degraded sequentially while a new DNA strand forms complementary to the transiently formed single-stranded DNA.

Stage II of the simulation approach models the formation of a single-stranded DNA from the hybrid, in what we can call ‘cleavage’ of the hybrid. An RNA-DNA bond is stochastically allowed to break once the bond length is shorter than a threshold (10 A ). This generates a single-stranded DNA (ssDNA), also called the complementary DNA (cDNA). It is important to note that in our model only one RNA chain undergoes this entire process, with the second strand staying inert. Strand transfer events involving the second RNA strand (3, 54, 55), which is known for introducing genetic diversity via RT, have been neglected here for simplicity.

Once the cDNA is formed, a second round of CG-KMC is carried out (Stage III) to simulate dNTP addition to cDNA, finally forming the double-stranded DNA. We note that the growth of dsDNA is initiated at the two polypurine tracts (PPT): the 3′-PPT and the central PPT (cPPT) (36, 62–64) (**Figure S8**). These are regions which are resistant to the nucleolytic degradation activity of RT-associated RNase H. The protocol for modeling this simultaneous growth of dsDNA from the two primers is discussed in detail in the SI. However, owing to structural and kinetic complexity, the three-stranded central DNA flap (62) was not explicitly modeled. Higher force constants were used for representing the newly formed bonds and angles, to accurately incorporate the stiffness and rigidity of dsDNA. This rigidity plays a crucial role in the uncoating of the capsid. In Stage III as well, an experimentally motivated “switch” is activated at ∼3.5 kb of the genome size (32) to initiate uncoating. This mimics the ultimate failure of the condensing effect of the NC protein (discussed in the Results section) and implicitly recapitulates the conditions that drive RT. A total of five sets of simulations were carried out for Stages I and II. For each set in Stage III, three runs were performed, each corresponding to a different ε value in Eq (1).

### CG-MD simulations, data analysis, and visualization

All CG-MD simulations were carried out using LAMMPS (21 July 2020) (65). The equations of motion were integrated with the Velocity Verlet algorithm, and the system was periodic in all three dimensions. CG-KMC was performed through an iterative series of short MD simulations, each consisting of 10^4^ CG time steps, and each such iteration was carried out for each RNA/DNA bead undergoing bond formation/breaking. The CG timestep used was 50 fs (but it must be noted the CG “time” cannot be easily associated with a real time – the CG time is much longer than all-atom time). A Langevin thermostat (66) was used for propagating the nuclei at 310 K. The simulation box taken was 1500 A in all three spatial dimensions. Each short CG simulation allows exactly one CG-KMC move, which may be either bond making or breaking, at 50% probability. Frames were saved after every 10^4^ CG steps. The resulting trajectories and data were visualized with Visual Molecular Dynamics (VMD) (67). The analysis was performed using MDAnalysis (68), Gnuplot, and custom Python and Bash scripts. Green-Lagrange strain calculations were carried out with OVITO (69).

## Acknowledgements

Research reported in this publication was supported by the Behavior of HIV in Viral Environments (B-HIVE) Center of the National Institute of Allergy and Infectious Diseases (NIAID) of the National Institutes of Health (NIH) under award number U54AI170855 and through NIAID grant R01AI178850. The content is solely the responsibility of the authors and does not necessarily represent the official views of the National Institutes of Health. The authors also acknowledge the computational resources provided by the Research Computing Center (RCC) of The University of Chicago, Bridges-2 at the Pittsburgh Supercomputing Center through the Advanced Cyberinfrastructure Coordination Ecosystem: Services & Support (ACCESS) program, funded by the National Science Foundation grant numbers 2138259, 2138286, 2138307, 2137603, and 2138296, and Frontera at the Texas Advanced Computing Center (TACC), funded by the National Science Foundation (NSF grant OAC-1818253). K.G. acknowledges the John C. Light Memorial/John A. Weil Fellowship provided by the Department of Chemistry, The University of Chicago. K.G. also thanks Prof. Vinay Pathak, Dr. Arpa Hudait, Dr. Ryan Burdick and Dr. Curt Waltmann for valuable discussions.

## Data Availability Statement

All relevant codes and input scripts for performing CG-KMC and CG-MD simulations in LAMMPS will be made available on GitHub upon publication. Due to the large size of the files, trajectories will be provided upon request.

