## Supplementary Information for "Mechanism of HIV-1 Capsid Rupture and Uncoating by Reverse Transcription"

### Capsid starts uncoating as the double-stranded DNA grows

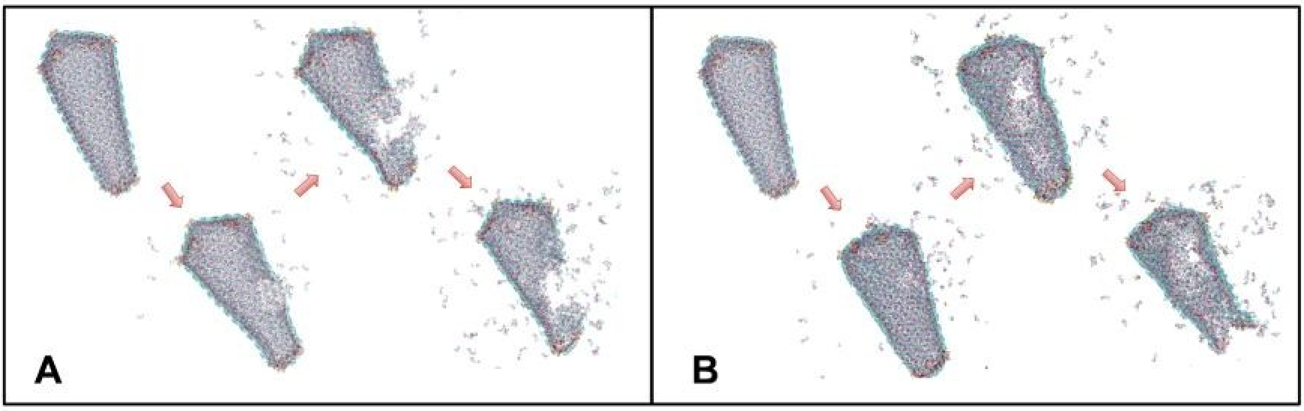


**Figure S1: Two representative pathways of capsid uncoating are illustrated: (A)** Rupturing, predominantly marked by detachment of hexamers from the midsection of the capsid lattice, progresses from the formation of small local defects to extensive lattice disassembly **(B)** An alternative pathway initiates with local defects near the wider end of the capsid and propagates longitudinally through the midsection toward the narrow tip, ultimately leading to significant lattice disruption.

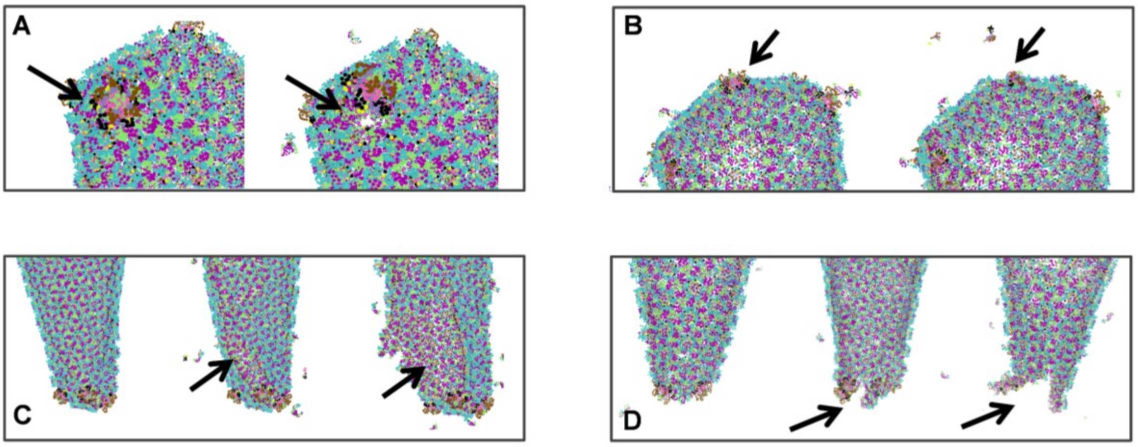


**Figure S2: Representative zoomed-in snapshots of lattice rupturing: (A)** A close-up of the wider end of the capsid shell, characterized by a gradual detachment or 'opening' of a pentamer (brown-black moiety). An 'empty' space is evident, as the pentamer gets detached. **(B)** A similar, but less prominent, pentamer detachment from the top of the capsid. **(C)** Longitudinal rupturing, initiated in the midsection, but gradually progressing towards the tip. **(D)** Prominent rupturing at the narrow end of the capsid.

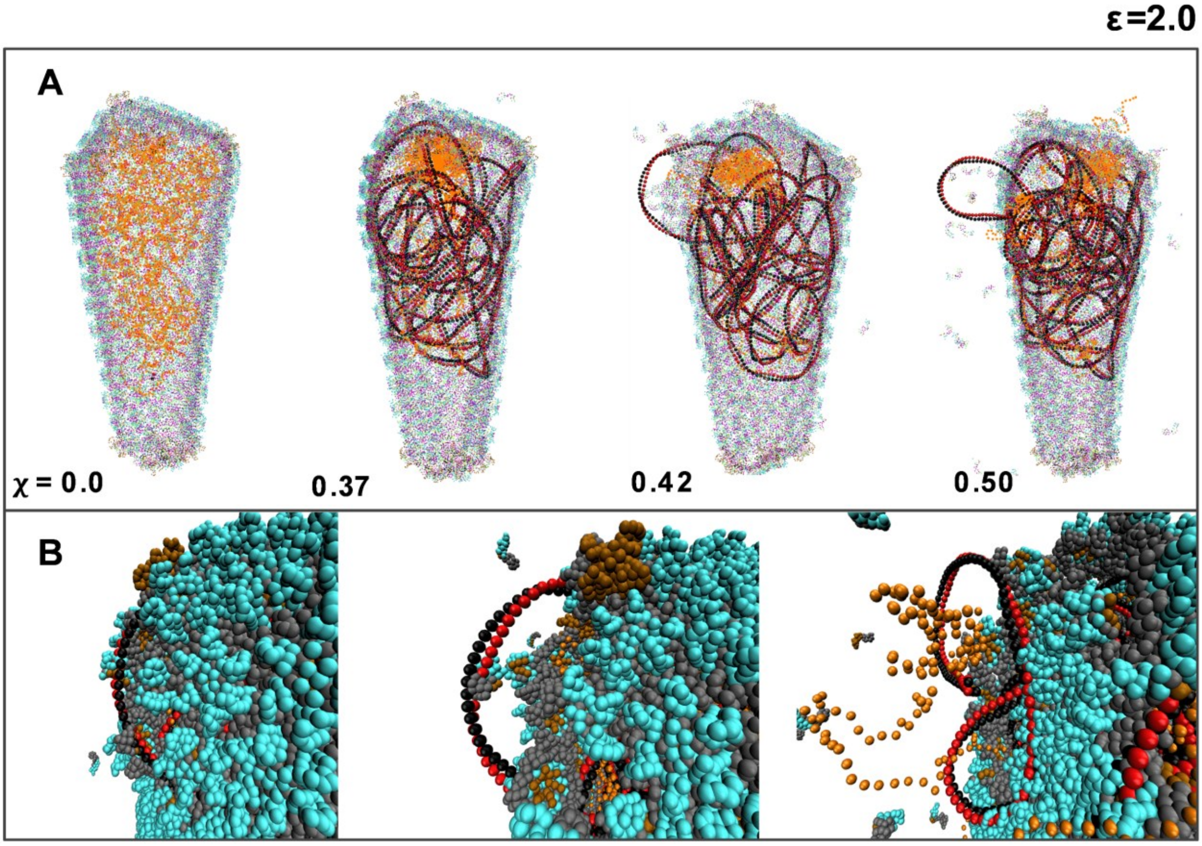


**Figure S3: For ε = 2.0 kcal/mol, the growth of dsDNA is depicted: (A)** Snapshots of the capsid enclosing the genome are shown. Initially, the genome is the cDNA (orange). CG-KMC is carried out on the cDNA to form dsDNA (black–red hybrid). As dsDNA grows, rupturing occurs near the wider end of the capsid, and the dsDNA eventually loops out prominently. **(B)** A closer view of the uncoating process. Clear signs of dsDNA protrusion can be seen, accompanied by detachment of both capsid pentamers and hexamers from the lattice. As RT progresses, the capsid walls begin to disassemble, releasing loops of dsDNA. Strands of cDNA also appear to loop out. Only one representative pathway for ε = 2.0 kcal/mol is shown. Here, χ represents the fraction of the total genome size, and the approximate CG timestamp corresponding to a configuration can be given as $\chi \times 3000 \times 10^4$ CG steps.

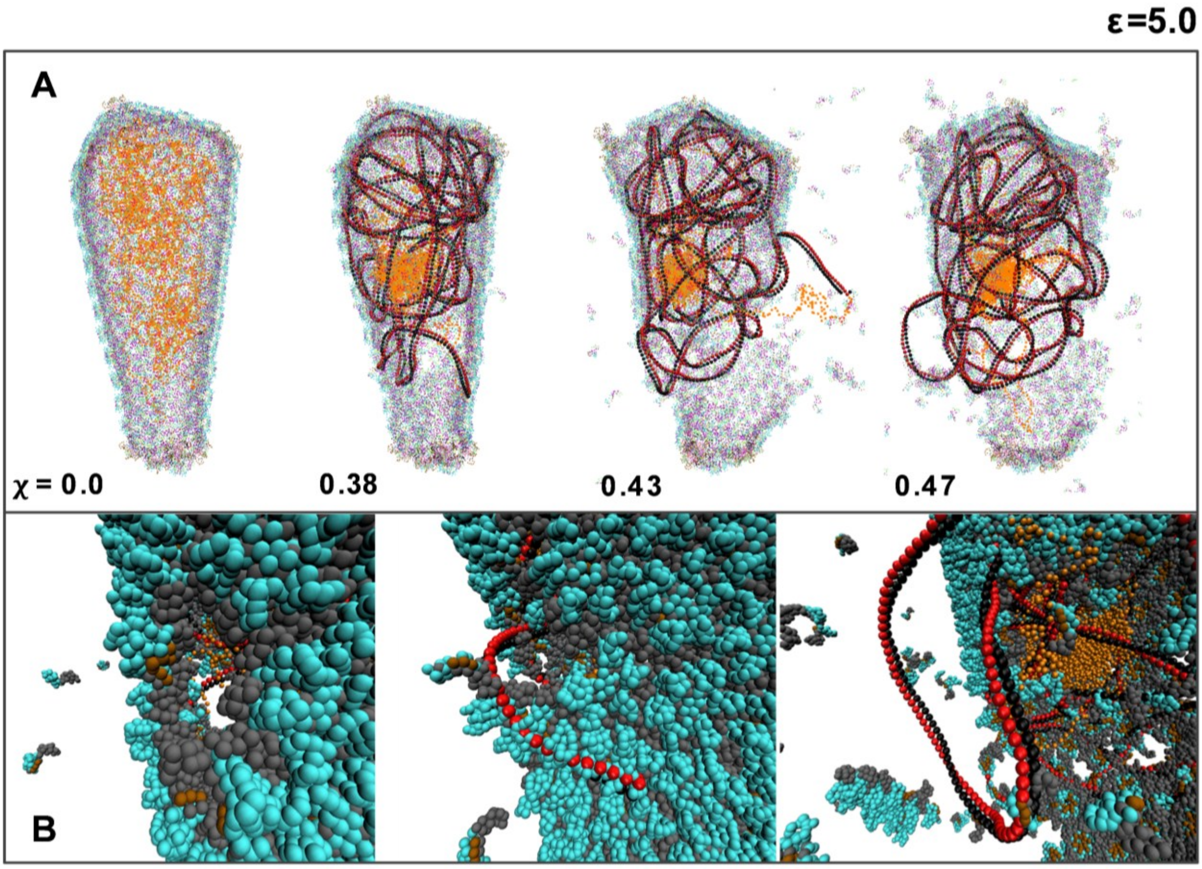


**Figure S4: For ε = 5.0 kcal/mol, the growth of dsDNA is shown: (A)** Snapshots of the capsid enclosing the genome are presented. Initially, the genome consists of cDNA (orange). CG-KMC is carried out on the cDNA to form dsDNA (black–red hybrid). As dsDNA grows, severe rupturing occurs near the midsection of the capsid, ultimately leading to partial disassembly, with clear dsDNA protrusions visible. **(B)** A closer view of the uncoating process shows distinct dsDNA protrusion accompanied by pronounced uncoating as the strands loop outward. Uncoating is markedly more substantial than for ε = 2.0 and 3.5 kcal/mol. Only one representative pathway for ε = 5.0 kcal/mol is shown. Here, χ represents the fraction of the total genome size, and the approximate CG timestamp corresponding to a configuration can be given as $\chi \times 3000 \times 10^4$ CG steps.

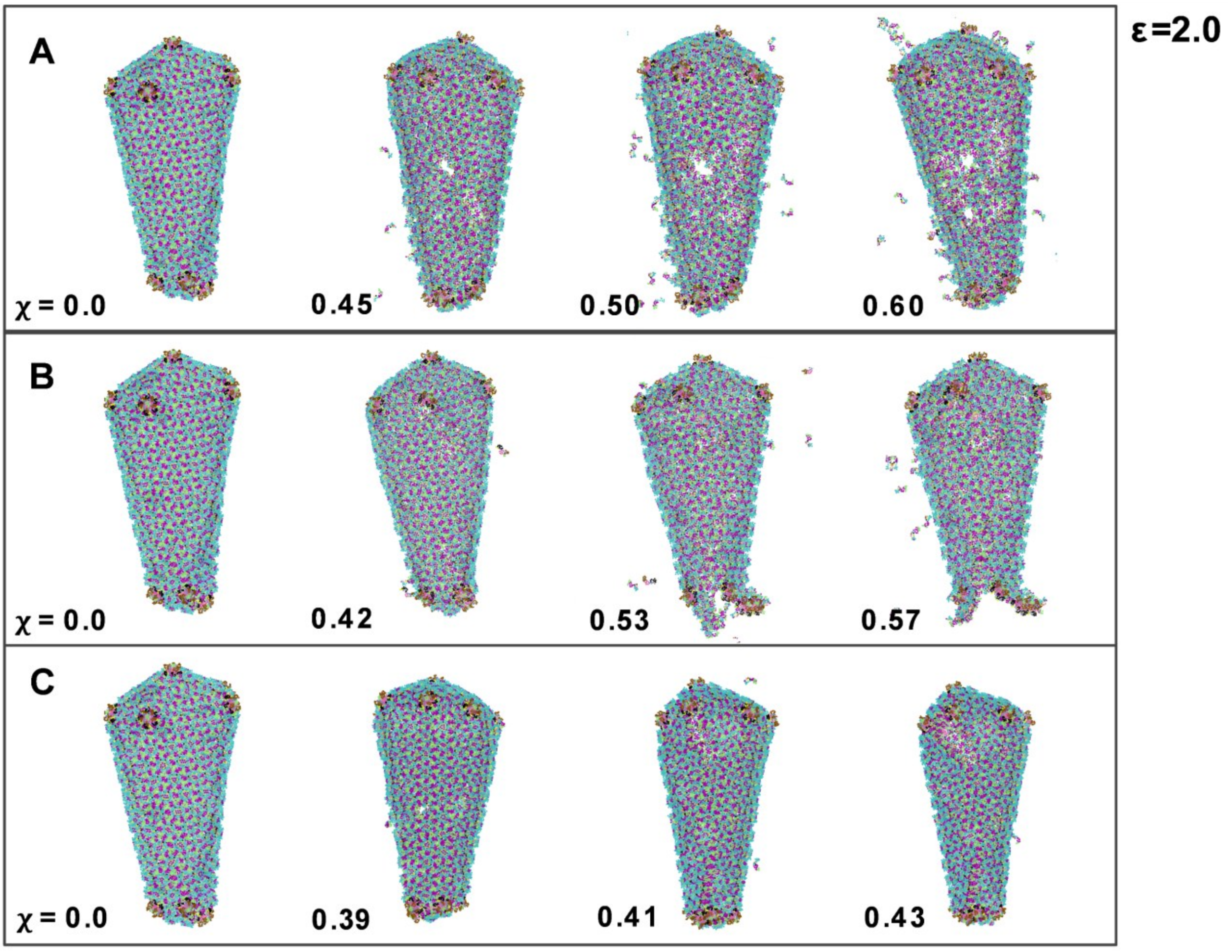


**Figure S5: Three representative capsid uncoating pathways for ε = 2.0 kcal/mol: (A)** Small cracks first emerge in the midsection of the capsid lattice as RT proceeds, gradually expanding into larger openings. **(B)** Cracks initiate at the narrow end of the cone-shaped capsid and are marked by the detachment of both pentamers and hexamers. **(C)** Very slow, gradual crack formation occurs near the wider end of the capsid, with rupture propagating along these cracks toward the midsection of the lattice. Here, χ represents the fraction of the total genome size, and the approximate CG timestamp corresponding to a configuration can be given as $\chi \times 3000 \times 10^4$ CG steps.

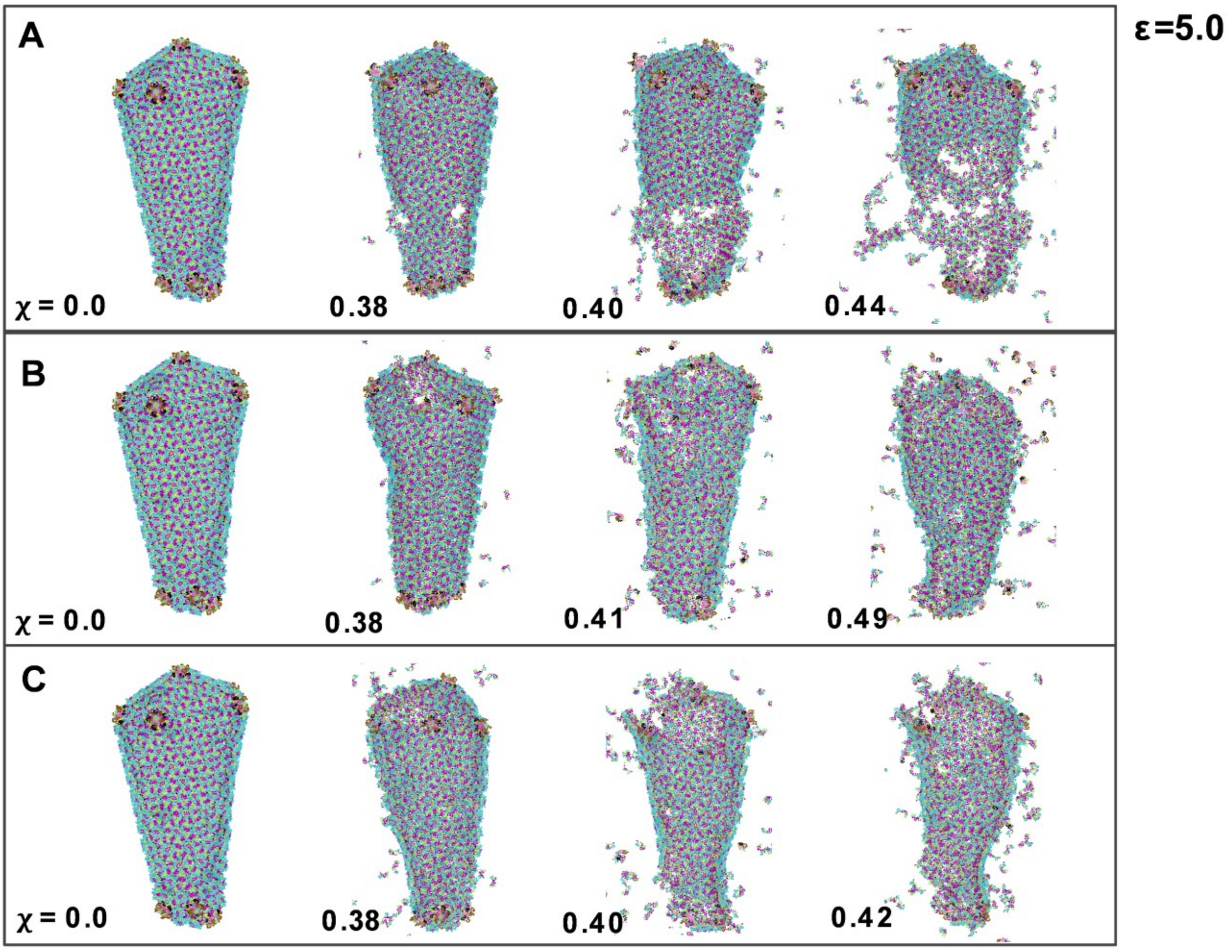


**Figure S6: Three representative capsid uncoating pathways for ε = 5.0 kcal/mol: (A)** Cracks originating in the midsection of the lattice eventually tear apart the capsid, resulting in a crumbling, heavily ruptured structure. **(B)** Distortions and cracks form openings interspersed along the capsid walls, with the most prominent ones appearing near the wider end. **(C)** Cracks emerging near the wider end of the lattice gradually widen and propagate along the vertical axis, ultimately causing extensive rupturing of the entire capsid. Here, $\chi$ represents the fraction of the total genome size, and the approximate CG timestamp corresponding to a configuration can be given as $\chi \times 3000 \times 10^4$ CG steps.

### Green-Lagrange strain calculations at CG resolution

To quantify the local deformation of a CG bead with respect to its equilibrium position (in this case, a stable capsid lattice before uncoating), the Green-Lagrange strain tensors were computed. The per-particle Green-Lagrange strain tensor is defined as:

$$\mathbf{E} = {}^{1}\!/_{2}\,(\mathbf{F}^{\mathbf{T}}\mathbf{F} - \mathbf{I}) \tag{S1}$$

where **F** is the local deformation tensor, computed from the particle displacement vectors in the reference and current configurations. The strain patterns for a representative uncoating pathway are depicted in **Figure S7**.

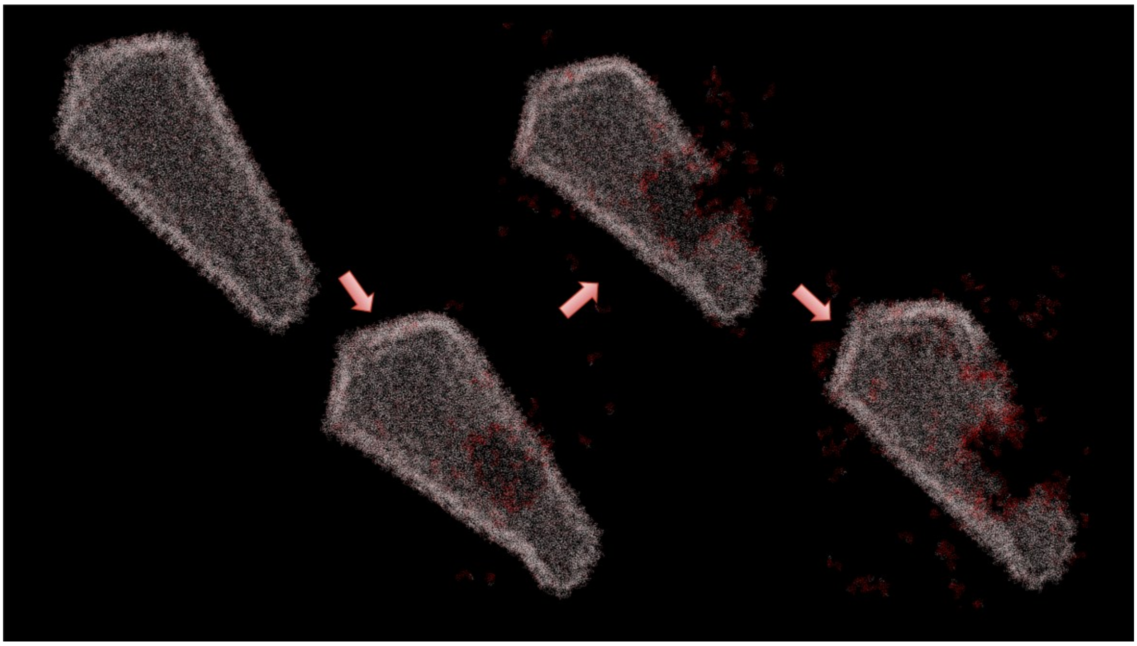

**Figure S7:** Green-Lagrange strain calculated for a representative uncoating pathway, characterized predominantly by rupturing at the capsid midsection. Regions of high particle strain, indicated by progressively deeper red hues (white indicating equilibrium position), localize primarily at the midsection and lead to rupture events. The most highly strained regions are concentrated along the periphery of the rupture site, highlighting the lattice boundaries most susceptible to mechanical failure.

### dsDNA synthesis is initiated at the polypurine tracts (PPT)

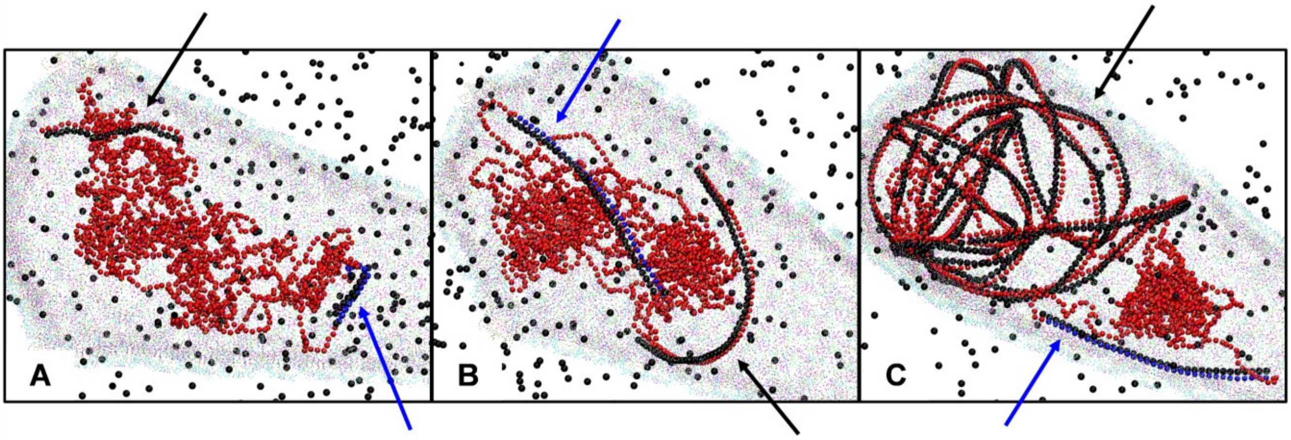


**Figure S8: dsDNA synthesis from cDNA initiated at the 3′-PPT and cPPT: (A)** Simultaneous growth of dsDNA from two distinct polypurine tracts (PPT): the 3′-PPT and the central PPT (cPPT). **(B)** Continued growth and elongation of both dsDNA strands. **(C)** The dsDNA initiated from cPPT spans 99 nucleotides (~33 CG beads), whereas the dsDNA initiated from 3′-PPT continues to elongate. In all panels, the blue-black hybrid (indicated by blue arrows) represents the dsDNA growing from the shorter cPPT, while the red-black hybrid (indicated by black arrows) represents the dsDNA growing from the 3′-PPT. The randomly distributed black spheres represent dNTP beads.

The dsDNA is synthesized from cDNA as two half-genomic strands (Ref. 59 of the main text) and is initiated at two regions that act as primers: the upstream plus strand from the 3′-PPT and the downstream plus strand from the cPPT. To approximately model the length and position of these segments, CG-KMC is initiated at bead index ~100 (or ~2900 due to symmetry in the CG model) of the cDNA to mimic 3′-PPT initiated dsDNA synthesis, and at roughly half the length of the genome (~1500 CG index) for the cPPT region. In accordance with experimental results (Refs. 59 and 60 of the main text), dsDNA growth from the cPPT was capped at 33 CG beads, corresponding to 99 nucleotides. However, the relative kinetics of dsDNA growth from the two primers with respect to each other is not well characterized. To model this, let $\xi \sim \mathcal{U}\ (0,1)$ be a random number drawn from a uniform distribution between 0 and 1. Now, we define a criterion ($C$) to determine, at a given MD step, which primer/region participates in dsDNA growth:

$$C = \mathbf{\Theta}(\alpha - \xi) \tag{S2}$$

where, the Heaviside function is given by

$$\mathbf{\Theta}(x) = \begin{cases} 1 & x > 0 \\ 0 & x \leq 0 \end{cases} \tag{S3}$$

If $\xi < \alpha$, dsDNA evolves from 3′-PPT, else from cPPT. As 3′-PPT is predominantly responsible for dsDNA growth, α was set to 0.7, 0.8 and 0.9 to reflect 70-90% probability of 3′-PPT associated dsDNA growth.

Representative snapshots depicting the explicit growth of two dsDNA segments from both primers are shown in **Figure S8**. As is evident there, the growth of the longer DNA strand (corresponding to 3′-PPT) is accompanied by the simultaneous growth of another strand (corresponding to cPPT). Replicas with different α values were carried out to reflect different relative kinetics in the growth of the two dsDNA strands with respect to each other. It is critically important to note that the capsid rupture patterns are not influenced by the mode of dsDNA growth, whether it occurs continuously from 3′-PPT alone or in a segmented manner from both PPT regions. Extensive tests and replicas performed with and without the cPPT region demonstrate that capsid rupture primarily depends on the overall fraction of the viral genome that has been reverse transcribed, that is, on the total bulk of dsDNA formed.

### Capsid rupture patterns driven by a model virtual object expansion

It is tempting to assume that the rupture of the capsid by RT occurring within it can be modeled by some "virtual object" inside that exerts outward pressure. This turns out to be a flawed assumption. To demonstrate this, we also investigated the capsid rupture patterns not as a consequence of an explicit RT model as in the rest of this paper, but as a consequence of a virtual object expanding inside the capsid. We thus embedded a virtual object inside the capsid, with a pre-defined geometry (**Figure S9**). As this object isotropically grows inside the capsid, it exerts uniform repulsive forces on the proteins outside the object (in this case, all capsid CG beads are outside the virtual object), effectively pushing the CG particles outward. Once a particular threshold is exceeded, the capsid walls begin to rupture. **Figure S9** demonstrates two cases: (A-B) expansion of a sphere (C-D) expansion of two spheres, stacked on top of each other, inside the capsid. We call the latter a "double sphere" object. A sphere, initially placed near the wider end of the capsid, has an initial radius ($R_{s,init}$) of 5 Å. The growth equation used was:

$$R_{s,new} = R_{s,init} + \mathbf{\Delta}R \cdot n_{\boldsymbol{steps}} \tag{S4}$$

where, $\Delta R$ = 0.001 Å and $n_{steps}$ refers to the MD simulation steps.

In the case of the ”double sphere” object (which might be argued to be a more realistic capsid contents), a smaller sphere is stacked on top of a larger one, such that the radius of the larger one is maintained at twice that of the smaller sphere throughout. To preserve this ratio, a growth rate of $\Delta R$ = 0.002 Å was used for the larger sphere, with an initial radius of 10 Å. The smaller sphere ($\Delta R$ = 0.001 Å with an initial radius of 5 Å) was placed near the wider tip of the capsid.

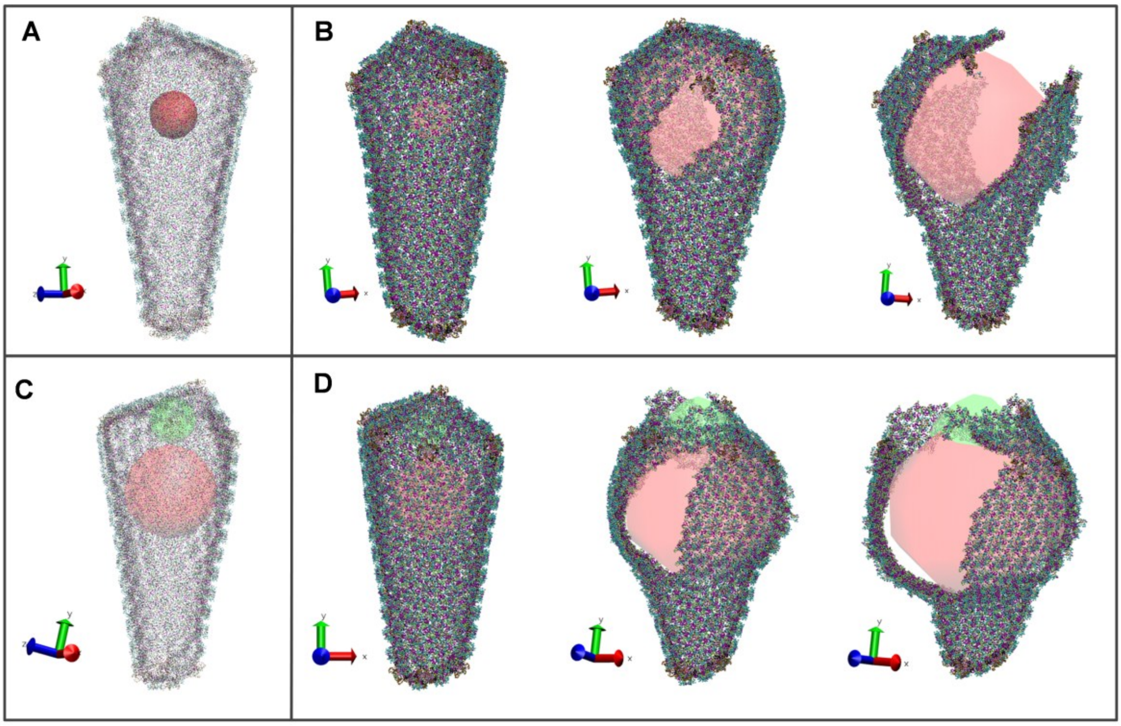


**Figure S9: Capsid rupture as a function of a dynamically expanding virtual object within the capsid: (A)** A red virtual sphere is embedded inside the capsid near the wider end. **(B)** The capsid rupture pathway occurs as the sphere grows, eventually causing the capsid to tear near the wider cone. **(C)** Two virtual spheres are embedded inside the capsid with the smaller one (green) on top of the larger one (red). **(D)** The capsid rupture pathway occurs as the spheres expand (green axis), eventually detaching the wider tip of the capsid from the rest of the lattice.

**Figure S9** shows distinct capsid rupture pathways as the virtual object grows. As the virtual sphere expands (**Figures S9A-B**), the wider end of the capsid starts bulging out, leading to a relatively unphysical deformation. With continued sphere expansion, a local defect emerges along the capsid wall and spreads diagonally, resulting in extensive rupture that gives the appearance of the capsid ‘opening up’. On the other hand, as the double sphere expands (**Figures S9C-D**), a longitudinal crack appears near the wider end and gradually propagates, in parallel with rupture at the wider tip. As both the spheres expand, the capsid once again appears to ‘open up’, with an additional patch forming at the tip. Unlike the RT driven capsid rupture simulations which are predominantly stochastic in nature leading to multiple anisotropic rupture patterns, the virtual object expansion driven rupture patterns are rather deterministic, i.e., expansion of the same geometric object leads to identical rupture patterns across multiple replicas. The rupture patterns in **Figure S9** were carried out over three replicas, for each virtual object, and demonstrated virtually the same rupture pathways. Moreover, the virtual object expansion scheme does not generate very realistic capsid ruptures that agree with experimental cryo-ET images (**Figure 4** of the main text), unlike the RT approach. As this is rather deterministic, capsid rupture patterns can be altered by tuning the geometric parameters of the virtual object, but none of them we looked at were realistic. This has been extensively discussed in the main text – significantly demonstrating the need for explicitly modeling RT (as in this work) that can drive anisotropic capsid rupture, as a direct result of dsDNA growth. In essence, the dsDNA as it grows acts a bit like a rather rigid and growing "snake" that can dynamically seek out vulnerabilities in the capsid lattice, especially in high curvature regions, and rupture them.

### Mathematical formulation of CG-KMC

Let $\mathbf{r}_i, \mathbf{r}_j \in \mathbb{R}^3$ denote the positions of two CG particles. Let $r_{cut}$ be the distance cutoff for the CG-KMC move. Let $\xi \sim \mathcal{U}\ (0,1)$ be a random number drawn from a uniform distribution between 0 and 1. Now,

$$r_{ij} = ||\, \mathbf{r}_i - \mathbf{r}_j\, || \tag{S5}$$

Therefore, the CG-KMC move acceptance, at a given MD step, can be denoted as:

$$A = \mathbf{\Theta}(r_{cut} - r_{ij})\mathbf{\Theta}(0.5 - \xi) \tag{S6}$$

where, $\mathbf{\Theta}(x)$ is once again the Heaviside function (as in Eq (S3)).

Thus, a CG-KMC move is accepted only if $r_{ij} < r_{cut}$ and $\xi < 0.5$, to reflect 50% probability of acceptance. For Stages I and III in RT, $r_{cut} = 20$ Å, and 10 Å for Stage II. In Stage I, a bond forms between a CG RNA bead and an incoming dNTP bead. In Stage II, the bond in the resulting RNA–DNA hybrid is cleaved (thus, $r_{ij} > r_{cut}$), thus producing cDNA. Finally, in Stage III, a bond forms between the newly formed cDNA and a dNTP, completing the sequence of events. We note that the use of the word "kinetic" here in the algorithm title means that kinetic (reactive) processes are associated with the MC moves. In typical KMC algorithms for simpler systems (e.g., an adsorbate atom diffusing on a corrugated metal surface), one can also associate a timescale with each KMC move and then update the clock so that the KMC algorithm becomes temporal (dynamic in the true sense of the word). We do not do this in the present work because the RT process is more complicated than a single (or few) barrier crossing features and the timescales of the elementary RT steps are not known in detail anyway.

**Movie S1:** Two representative pathways of capsid rupture, including the dynamic growth of dsDNA from ssRNA. Pathway **A** depicts capsid rupture near the wider end of the cone, whereas pathway **B** depicts severe rupture across the capsid walls near the narrower end.
